# The geometric structure of features underlies human VTC object recognition

**DOI:** 10.64898/2026.01.22.696490

**Authors:** Bincheng Wen, Chuncheng Zhang, Changde Du, Le Chang, Huiguang He

## Abstract

The ventral temporal cortex (VTC) plays a crucial role in human object recognition. Within VTC neural space, object-specific domains align with domain-general features, e.g. face- and scene-domain align with the feature distribution of animacy, generating a hierarchical knowledge structure for categorization. However, the neural process of integrating information from VTC to distinguish different objects remains unclear. Here, we employed a combination of ANN modeling, functional MRI, and MEG to investigate how VTC features affect object manifold separability. The representational geometry analysis shows that domain-general features in VTC form a unique structure, different from ANN, to assist object classification. Moreover, VTC dynamically adjusts the geometrical relationship of these features during object recognition, influencing the geometrical properties of object manifolds and, consequently, their separability. These findings advance our understanding of the neural computation involved in VTC object recognition, revealing how downstream neurons can flexibly access category information in diverse recognition tasks.

## Introduction

Primates possess a remarkable ability to rapidly identify target objects in natural environments. This ability is largely attributed to the ventral visual stream (VVS) (Ungerleider, 1982), a complex neural process that spans multiple areas of the cortex, transforming the optical signal from the retina into category-related responses within the ventral temporal cortex (VTC). Within the VTC, neurons with distinct category selectivity concentrate in different cortical areas (Aparicio et al., 2016), each specializing in recognizing a particular object domain, such as faces (Kanwisher et al., 1997) or places (Epstein and Kanwisher, 1998). These domain-specific areas are anatomically arranged along domain-general features (Bao et al., 2020; Kriegeskorte et al., 2008), such as animacy (Sha et al., 2015; Wiggett et al., 2009) and real-world size (Konkle and Oliva, 2012), creating a hierarchical knowledge framework encoded by VTC that aligns with subjects’ behavioral judgments (Grill-Spector and Weiner, 2014; Rosch et al., 1976). However, the neural computational mechanism of how VTC effectively integrates domain-general features to distinguish one object from another remains a key open question for cognitive and neural scientists.

One promising hypothesis, building on the functional topography of domain-general features, is that VTC constructs a general object space to describe perceived objects (Bao et al., 2020; Yao et al., 2023). This scheme suggests that objects perceived as similar at the category level are grouped, while dissimilar ones are separated, resulting in various low-dimensional manifolds — object identity manifolds — in the high-dimensional VTC neural space (DiCarlo and Cox, 2007). Separating neural manifolds allows a putative downstream neuron to determine an object’s identity using a linear hyperplane based on the pattern of VTC responses it elicited (DiCarlo et al., 2012; Pagan et al., 2013). Additionally, the linear separability within the object space offers great behavioral flexibility by maximizing the potential for discrimination among diverse objects and their combinations (Grill-Spector and Weiner, 2014). Yet, it remains unclear how the VTC organizes domain-general features into object spaces to maximize object manifold capabilities.

Recent research suggests that the separability of object manifolds is determined by their geometrical properties (Cohen et al., 2020; SueYeon, 2017). Indeed, within a neural space, separating object manifolds becomes simpler as their distance grows, the asymmetry of manifold size results in the classification hyperplane’s bias towards the larger manifold, and the manifolds’ dimensionality reduces the noise impacting their separation (Sorscher et al., 2022). However, it’s still unclear how domain-general features affect the separability of object manifolds. Furthermore, during the process of object recognition, VTC’s representation of features exhibits dynamic changes over time (Cichy et al., 2014; Kietzmann et al., 2019), leading to potential changes in the geometrical relationship of the features. One intriguing possibility is that the geometrical structure of VTC features dynamically constrains the object manifolds, providing a potential mechanism for inferring the identity of a single object. As a result, the dynamic geometry structure of domain-general features serves as structural knowledge represented in the VTC, facilitating putative downstream neurons to flexibly obtain key information for distinguishing objects.

In this study, we first investigated the geometry relationship between domain-general features in VTC. We then hypothesized that if this geometric relationship is important for inferring object identity, it would affect the manifold separability of objects by constraining the manifold geometry. Building on this, we further proposed that VTC would dynamically modulate the feature geometry based on task requirements.

To test these hypotheses, we first mapped three predominant domain-general features onto the VTC through the elicited fMRI responses when human participants passively viewed the representative images. Subsequently, we explored the features’ geometric structure in the VTC neural space. Relative to the ANN model, the observed feature geometry of VTC constrained the object manifolds with a divergence feature structure, providing potential advantages for recognizing specific objects. By combining computational models with MEG datasets, we found evidence that VTC dynamically modulates the responses of domain-general features, facilitating the separation of the objects that were recognized. Our findings extend the conventional understanding of the neural processing of object recognition in the ventral temporal cortex, indicating that the geometrical structure of domain-general features, as structural knowledge encoded by VTC, underlies the flexible object recognition capabilities of humans.

## Results

### Mapping domain-general features onto VTC

We first independently mapped the domain-general feature to the VTC one by one to minimize the bias caused by the cognition of specific objects (Çukur et al., 2013; Kay and Yeatman, 2017). To capture the domain-general features in the VTC, we leveraged the well-studied artificial neural network (ANN), Alexnet (Krizhevsky et al., 2012), which provided an approximate framework for the feedforward process of the ventral visual pathway (Cadieu et al., 2014). From the 200k ImageNet dataset (Deng et al., 2009), we extracted candidate features of the VTC and identified the top three main features for object recognition through principal component analysis (PCA) (see Methods, Fig.1a∼c). These three features were related to the well-known VTC representations: animacy (Sha et al., 2015), curvature (Yue et al., 2014), and real-world size (Konkle and Oliva, 2012) (Supplementary Fig.2a∼c). A single node on these feature axes is often associated with a specific object category, causing potential bias in the mapping process. To address this, we carefully selected four nodes on each feature axis and utilized multiple representative images of these nodes as the stimuli for the fMRI experiment.

**Figure. 1.**
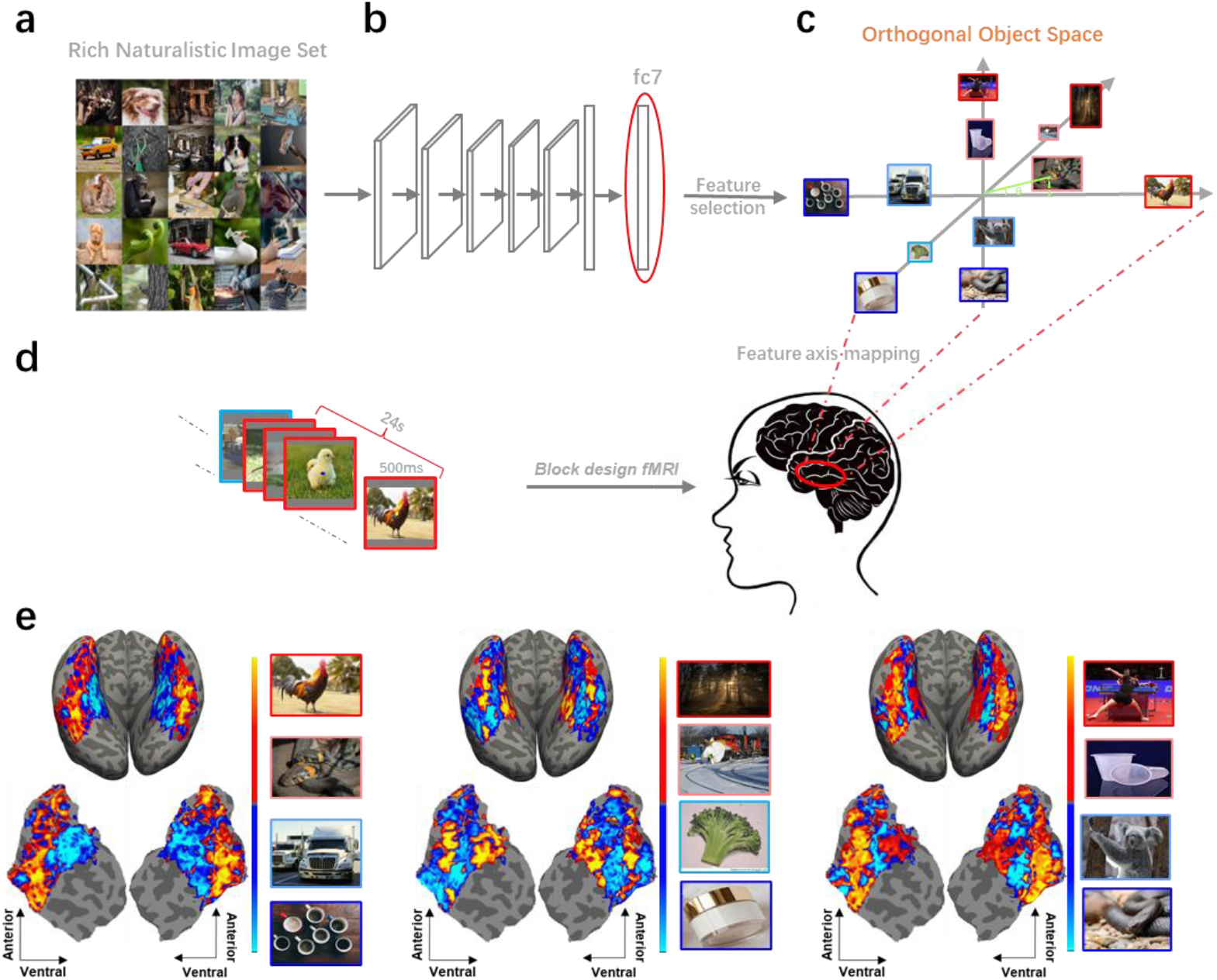
Domain-general feature mapping. **[Note: Due to copyright reasons, the images displayed have been replaced with similar images from Pixabay and are under a Creative Commons licence CC BY 0.]** **a**. This study employed a diverse naturalistic image set, encompassing 200 pictures for each of the 1000 categories, resulting in a total of 200k images. These images were randomly selected from the ImageNet 2012 train set. **b**. Domain-general features were distilled from the penultimate layer of Alexnet through PCA. **c**. The object space formed by these features was used to select feature nodes. The angle between the image and the feature axis is represented by a green solid line, while the image projection position on the axis is represented by a dotted line. The color of the image frame corresponds to the different nodes along the feature, consistent in **d. d**. Individual features were mapped to VTC through block design fMRI experiments. Each feature node block contained 48 images from the same node, each image presented in random order for 500ms. **e**. The VTC topography of the domain-general feature of the subject 04. The main distribution direction of the feature nodes was regarded as the feature direction. Colors on the brain surface indicate the preference of VTC voxels for features. Due to copyright reasons, the images displayed have been replaced with images from Pixabay and are under a Creative Commons licence CC BY 0.

We conducted a block-design fMRI experiment to collect responses for feature nodes. Positive and negative feature node blocks of the same feature were alternately presented in each run, with participants passively viewing representative images on the screen. Additionally, a control block of natural images from the ImageNet dataset, flanked by gray-screen blocks, was included in the middle of each run (Supplementary Fig. 3a). Out of the nine participants scanned, five completed all the fMRI experiments and were included in the following analysis. The remaining four were excluded due to incomplete participation. For the five subjects, each feature node was repeated 32 times across multiple scanning sessions (∼8 h scanning in each of N = 5 subjects), resulting in highly reliable responses (Supplementary Fig. 4).

Our feature mapping experiment enables us to obtain the single feature representation by identifying the main distribution direction of the feature nodes in the VTC neural space. We evaluated the quality of mapping results from three aspects. First, we tested the consistency of the feature dimensions in the VTC and ANN using the participation ratio (PR) (Peiran et al., 2017). Dimensionality indicates the number of linearly independent features needed to describe the feature nodes. Our analysis revealed no significant difference between ANN and VTC in terms of dimensionality (two-tailed paired t-tests across hemispheres, dimensionality VTC ∼= ANN penultimate layer: t_10_ = -0.3947, p = 0.7022; Supplementary Fig. 5a), but we observed a significantly lower dimensionality in the early visual cortexes compared to ANN (one-tailed paired t-tests across hemispheres, the dimensionality of visual cortex < ANN penultimate layer, V1: t_10_ = -37.4574, p < 0.0001; V2: t_10_ = -29.8411, p < 0.0001; V3: t_10_ = -8.1882, p < 0.0001; V4: t_10_ = -4.4570, p < 0.01; Supplementary Fig. 5a). These results also suggested that the expansion of feature dimensions was also part of VVS processing for visual information. Secondly, to confirm the linearly independent features were the features we wanted to map, we examined the consistency of the distribution of feature nodes in VTC and ANN. The node distribution of each feature in ANN could well explain the distribution in VTC (adjusted R^2^, feature 01: 0.6684; feature 02: 0.7494; feature 03: 0.6986; Supplementary Fig. 5d), indicating that the nodes of the same feature were arranged along the same direction in VTC. Lastly, by projecting each domain-general feature onto the surface of VTC, we obtained three gap-free VTC feature maps, where topographies were consistent with previous studies. Figure 1e illustrates the VTC feature maps of subject 04 (others see Supplementary Fig.5e). Notably, the domain-general feature 01, related to animacy, elicited a positive response from the mid-fusiform sulcus (MFS) to the lateral VTC, and a negative response on the other side, consistent with the animacy gradient of VTC (Sha et al., 2015). Similarly, the topography of domain-general feature 02, related to curvature, exhibits similarities to the VTC representation of curvature described in (Yue et al., 2020). Taken together, these observations verified the validity of our feature selection and mapping process.

### Geometry of domain-general features in VTC

Next, we asked how the VTC integrates the three domain-general features. Since the features selected from ANN were shared with VTC, VTC might construct an orthogonal object feature space similar to ANN. To test this hypothesis, we employed representational similarity analysis to explore the neural geometry of feature nodes and compared the geometric relationships of features between the VTC and the penultimate layer of the ANN. While the VTC had the highest similarity with ANN among all the cortical regions of VVS, there was a significant gap in feature geometry between VTC and ANN (mean correlation with the RDM of 12 features nodes of ANN penultimate layer on ± s.e.m. across hemispheres and lower bound noise ceiling (NC), V1: 0.1540±0.0191, NC: 0.7769; V2: 0.1645±0.0348, NC: 0.7426; V3: 0.3311±0.0208, NC: 0.8917; V4: 0.3793±0.0169, NC: 0.7805; VTC: 0.4054±0.0103, NC: 0.8484; Fig. 2a), indicating that a special feature space was constructed to represent objects in VTC.

**Figure. 2.**
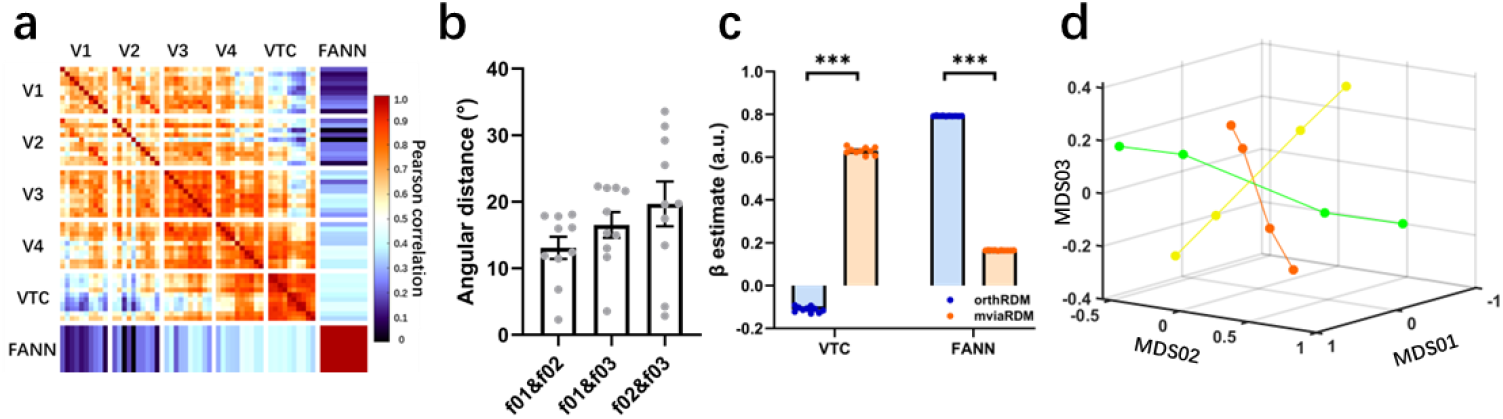
The geometry structure of domain-general features in VTC. **a**. Representation geometry similarity of the ventral visual cortices and ANN. Each small cell in the matrix represents the pairwise Pearson correlation of the RDM pattern of the 12 feature nodes between brain regions from different hemispheres or the ANN. 10 cells form large squares in the matrix, each corresponding to one brain area or the ANN penultimate layer. **b**. The absolute angular distance from the intersection angle of VTC features to the orthogonal angle. All the angle distances are away from zeros indicating that the three VTC features are not orthogonal with each other. Each data point represents the intersection angle of features within the VTC of one hemisphere, the error bars indicate s.e.m. **c**. Comparison of geometric model fits to VTC and ANN at the same noise level. orthRDM: the RDM of the model encoding orthogonal intersections, mviaRDM: the RDM of the model encoding the mean VTC intersection angles. Each data point represents the VTC of one hemisphere, or a noise-adding ANN penultimate layer, the error bars indicate s.e.m. **d**. The mean VTC geometric structure was reconstructed in a low-dimensional space. Each point is a feature node. Red, green, and yellow indicate domain-general features 01, 02, and 03, respectively. *p<0.05, **p<0.01, ***p<0.001.

To accurately quantify the feature geometry, we utilized representational similarity analysis (RSA) in conjunction with a parametric model-fitting approach (see Methods). Instead of fitting models encoding fixed geometric relationships between the three features, we explored the geometric relationship between VTC features by constructing representational dissimilarity matrices (RDMs) with continuously varying feature intersection angles. Our analysis confirmed that, in the VTC, domain-general features intersected with each other at non-orthogonal angles, distinct from the perpendicular intersections in the ANN (Angular distance to an orthogonal intersection (°) ± s.e.m. across VTCs: feature 01 & 02: 13.0766±1.6616; feature 01 & 03: 16.5203±1.9252; feature 02 & 03: 19.6983±3.3656; Fig. 2b). To estimate the potential impact of the noise in the BOLD signal on the divergence observed between VTC and ANN feature geometry, we conducted a noise-normalization procedure. Two model RDMs, one encoding orthogonal intersections (orthRDM) and the other capturing the mean VTC intersection angles (mviaRDM), were used to predict brain activity and ANN activity under similar noise conditions at the same time. The orthogonal model strongly correlated with ANN activity, while the VTC responses aligned best with the non-orthogonal model (one-tailed t-test across 10 VTCs, the weight of orthRDM > mviaRDM in ANN: t_10_ = 596.7862, p < 0.0001; the weight of mviaRDM > orthRDM in VTC: t_10_ = 79.2888, p < 0.0001; Fig. 2c) discounting noise as the source of divergence. The non-orthogonal intersection structure of domain-general features is illustrated in Fig 2d by Multidimensional Scaling (MDS) visualization of the best-fitting model RDMs for VTC in three dimensions.

### Lower-level visual cortices facilitate the formation of VTC feature structures

The geometric relationship of VTC features could reflect the static, stable structured knowledge of object categories encoded within VTC (Bugatus et al., 2017), but can also be modulated by the specific recognition task requirements from downstream brain areas (Çukur et al., 2013). To explore this further, we next investigated which brain regions, beyond local processing in VTC, might be involved in shaping the VTC feature structure. A surface-based searchlight RSA was used to assess whether the responses elicited by feature nodes were related to forming the VTC non-orthogonal intersection structure (see Methods; Fig. 3a). At the group level, VTC was a hotspot because its responses correlated well with its own feature structure. The lateral occipital-temporal cortex (LOTC) also showed a significant positive correlation with the VTC feature structure. Although subject 03 exhibited a slight negative correlation with the VTC feature structure in the lower-level visual cortices, this effect was not significant at the group level (Supplementary Fig. 6). Consistent with Fig 2a, the feature structure along the ventral visual pathway became more similar to the VTC feature structure. Beyond these ventral visual areas, there was no significant correlation between the feature structure of VTC and other brain regions at the group level, suggesting that the ventral visual processing was responsible for forming the feature structure we observed in VTC.

**Figure. 3.**
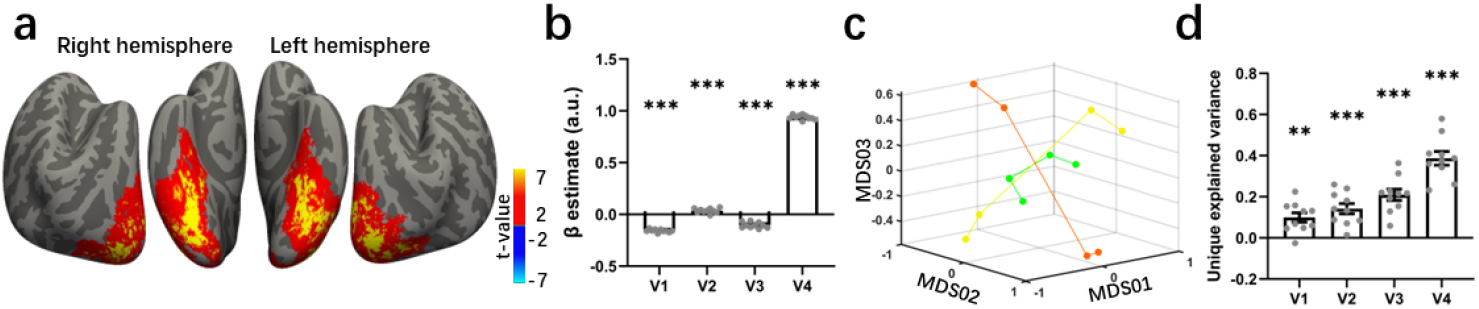
Basic VTC feature geometry arises from ventral visual processing. **a**. We tested which brain regions are related to the VTC feature geometry, using the surface-based searchlight RSA. At the group level, no significant connections were found in any brain regions, except for the ventral pathway. The data are masked to show only vertices where the correlation was significant (two-sided P < 0.05 via a permutation cluster corrected across subjects). **b**. Fitting the feature structure of VTC with the responses of four lower-level visual cortices: V1, V2, V3, and V4. Each data point represents an ROI of one hemisphere, and the error bars indicate s.e.m. **c**. The VTC feature structure predicted by the four lower-level visual cortices was reconstructed in a low-dimensional space. Each point is a feature node. Red, green, and yellow indicate domain-general features 01, 02, and 03, respectively. **d**. The unique explained variance of each ROI was calculated by comparing the explained variance between the model using both the ROI’s response and orthogonal feature structure to fit the VTC feature structure, with a model where the ROI’s response was replaced by a random variable. Each data point represents an ROI of one hemisphere, the error bars indicate s.e.m. *p<0.05, **p<0.01, ***p<0.001.

To test the role of visual cortices in shaping VTC feature structures, we used the responses of different regions of interest (ROIs) to predict the feature structure of VTC. The V1 and V3 showed a small but significant negative contribution compared with V2 and V4 (two-tailed group-level t-tests of the regression weights against zero: V1, t_10_ = -33.8592, p<0.001; V2, t_10_ = 4.8159, p<0.001; V3, t_10_ = -11.4210, p<0.001; V4, t_10_ = 128.4680, p<0.001; Fig. 3b). These results implied that compared to V2 and V4, V1 and V3 had a reverse effect on facilitating the formation of the VTC feature structure, resulting in VTC features being rotated into different directions. Based on this conclusion, along with the processing of VVS, the geometric relationship of domain-general features seems to have gone through a progressive adjustment to form a feature structure in VTC. For instance, a feature structure representing statistical relationships in the natural environment is captured in V1 and then progressively processed by the ventral visual processing pathway to generate a feature structure that more closely aligns with downstream cognitive needs in VTC. However, features of the high-level visual cortex VTC are hopelessly tangled with each other in lower-level visual cortices (DiCarlo et al., 2012). Thus, the direct contribution of the low-level visual cortex on the geometric relationship between VTC features needs to be treated with caution. Consistent with this view, the VTC feature structure predicted by the four lower-level visual cortices (Fig. 3c) was not fully disentangled compared to the mean VTC feature structure shown in Fig. 2d.

Next, we further examined whether the contribution from the lower-level visual cortices was the reason for the divergence in the feature structure between VTC and ANN. Compared to adding random variables, incorporating any single low-level visual cortex response into the process of fitting the VTC with ANN’s orthogonal structure can significantly enhance the explained variance of ANN to VTC’s (one-tailed signed-rank test across 10 brain hemispheres for improved explained variance>0: V1: p<0.01; V2: p<0.001; V3: p<0.001; V4: p<0.001; aIFS: p<0.01; pIFS: p<0.01; Fig. 3d). Moreover, within the VVS, a higher-level visual cortex contributed to more improvements in explained variance, suggesting a more significant role in shaping the VTC feature structure and providing further support to previous findings regarding the gradual adjustment of feature relationships.

Taken together, our findings suggest that the observed feature structure of VTC primarily arises from hierarchical processing within the ventral visual pathway, with minimal contribution from other brain regions. Different visual cortices play distinct roles in modulating feature geometric relationships, leading to the divergence of the feature structure between the VTC and ANN. This unique geometric relationship of VTC features is independent of specific object recognition and therefore may represent the basic knowledge of category characteristics, affecting the retrieval of object information in putative downstream neurons. Furthermore, based on this hypothesis, selectively enhancing relevant VTC features during the recognition of specific objects could modify the basic geometric relationship between domain-general features, facilitating the separation of object manifolds in a specific way to meet the recognition demands.

### Feature structures of VTC affect the separability of object manifolds

For putative downstream neurons, inferring the identity of an object requires the object manifold to exhibit linear separability in the object space defined by VTC features. Therefore, to test our hypothesis, we initially investigated how the feature structure of VTC influences the separation of objects manifolds.

First, we connected the separability of object manifolds to the geometric properties of their manifolds through the manifold signal-to-noise ratio (SNR). The manifold SNR reveals the relationship between manifold shape and separability by four interpretable geometric properties of manifolds: signal, bias, dimension, and signal–noise overlap (Sorscher et al., 2022). The signal represents the pairwise distance between the manifolds’ centroids, the bias describes the relative size of pairs of manifolds, the manifold dimension quantifies the number of dimensions along which the concept manifold varies significantly, and the signal–noise overlap quantifies the overlap between the signal direction and the manifold axes of variation. We trained a logistic regression classifier to measure the linear separability of four ecologically important categories, including faces, bodies, words, and places. The four categories could be linearly separated in the orthogonal space constructed by the three domain-general features, highlighting the crucial role of these features in object distinguishing (Fig. 4a). Consistent with the results of the linear classifier, the manifold geometry in the orthogonal space showed that different categories of manifolds, such as faces and scenes, were distant from each other, while similar subcategories, such as adult faces and baby faces, were close, enabling the distinction of categories, despite the asymmetrical distribution of the manifold SNR due to the classification advantage of larger manifolds (adjust R^2^ between manifold SNR and category classification accuracy, upper triangular: 0.8348; lower triangular: 0.8260, Fig. 4a, Supplementary Fig. 7).

**Figure. 4.**
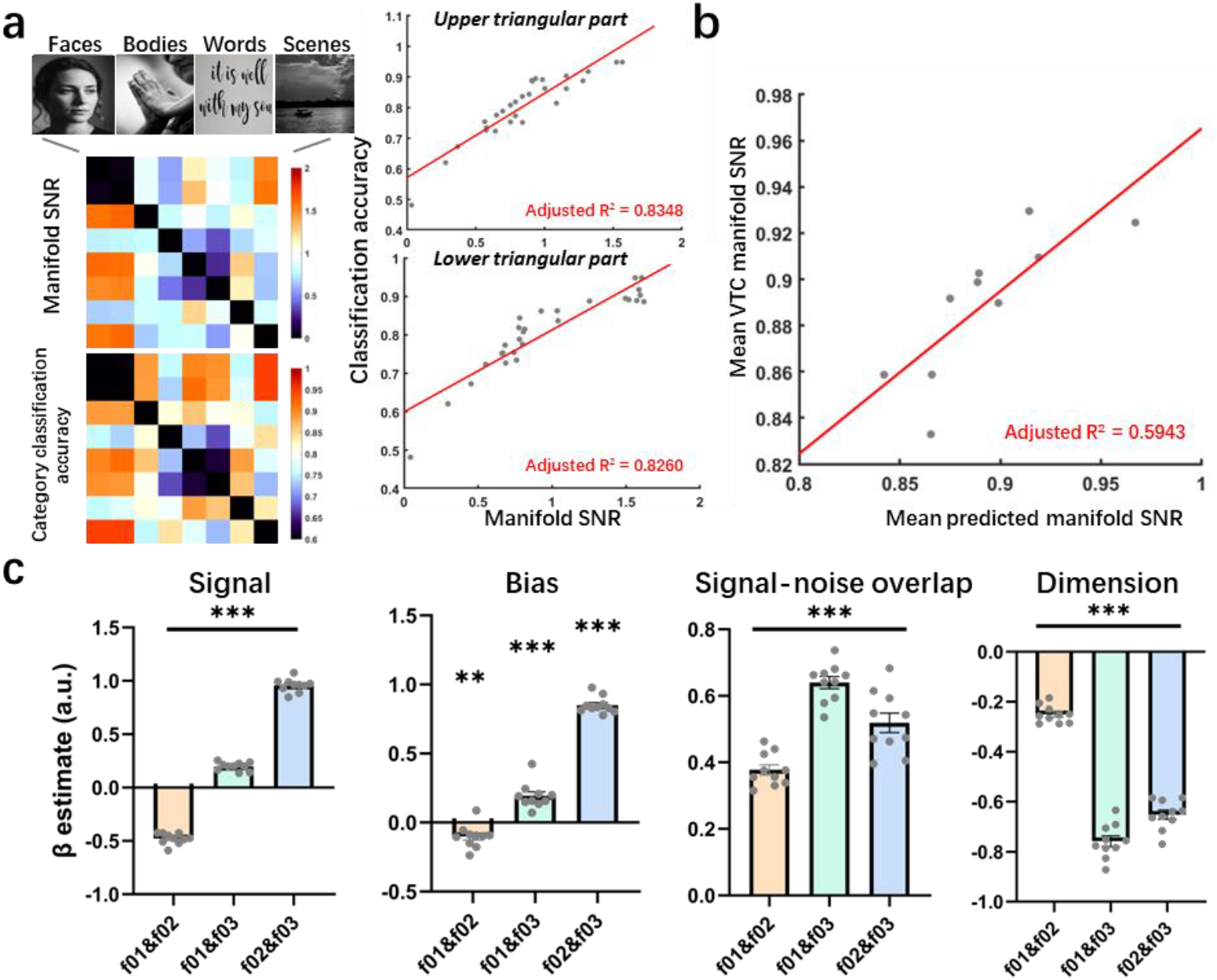
The feature structure of VTC constrains the geometry of object manifolds. **[Note: all human faces in this figure have been replaced by synthetically generated faces using the Nano Banana model (gemini-3-pro-image-preview) due to biorxiv policy on displaying human faces. Due to copyright reasons, the images displayed have been replaced with similar images from Pixabay and are under a Creative Commons licence CC BY 0.]** **a**. Connection between object discrimination and manifold shape in orthogonal object space. Four ecologically significant categories were used for analysis, each of which includes two subcategories: adult faces, baby faces, human trunks, limbs, numbers, letters, indoor scenes, and buildings. Left top, manifold signal-to-noise ratio between paired objects. Left bottom, the logistic regression classifier’s accuracy in discriminating paired objects. Right top and bottom, the correlation between the upper/lower triangular parts of the manifold SNR matrix and the matrix of classification accuracy. Each data point represents a paired relationship between two matrices. **b**. Top: A schematic diagram of how the feature rotation may affect the shape of object manifolds. Bottom: Correlation between the mean VTC manifold SNR and the mean manifold SNR predicted by the rotation angle of the three domain-general features. Each data point represents an experimentally measured basic VTC feature structure. The red line represents the result of linear fitting. **c**. Correlation between the rotated angle between two features and the four geometric properties that determine the manifold SNR. Each data point represents an experimentally measured basic VTC feature structure. The error bars indicate s.e.m. *p<0.05, **p<0.01, ***p<0.001.

Subsequently, we rotated the feature axis to the position of the measured basic VTC feature structure and evaluated the impact of this rotation on manifold SNR (see Methods). We found that the rotation angle of features could directly predict changes in manifold SNR (adjusted R^2^ between mean VTC manifold SNR and mean predicted manifold SNR: 0.5943, Fig. 4b), supporting the basic feature structure related to the VTC categorization process. Furthermore, we found that the rotation angle between features 02 and 03 was correlated with the average distance between manifolds, which promoted signals of separation between manifolds, while the rotation of features 01 and 02 had the opposite effect (Fig. 4c, signal). Similarly, the rotation angle between features 02 and 03 increased the average size of the manifold, providing a cognitive bias for distinguishing the manifolds, while the angle between features 01 and 02 had the opposite effect (Fig. 4c, bias). Additionally, the rotation of features reduced the average dimension of the manifold, deteriorating the signal-noise overlap between manifolds that were originally suppressed by high dimensions (Fig. 4c, signal-noise overlap and dimension). Therefore, feature structures comprehensively impact the geometric properties of manifolds, ultimately determining to what extent the manifold can be distinguished.

Together, our findings suggest that the feature structure of VTC can impose strong constraints on the geometric properties of neural manifolds, introduce cognitive biases in manifold separation, and achieve effective recognition.

### Domain-general features undergo a rotational transformation in object recognition

Next, we explored the role of feature structures in object recognition tasks. Our hypothesis suggested that the enhancement of relevant features during the recognition process could lead to geometric transformations between the features that constrain the neural manifold’s geometry, thus facilitating object discrimination. To confirm this inference, we additionally utilized a dataset that captured dynamic neural responses through MEG and fMRI as participants passively viewed 156 naturalistic images (Mohsenzadeh et al., 2019). At each time point, a Support Vector Machine (SVM) classifier was deployed on MEG sensor data to quantify the dissimilarity between image pairs, capturing the representational geometry of these images in neural space. Subsequently, the dynamic geometric relationship among the three domain-general features was explored using a parameterized RDM fitted with the neural RDM at each time point.

Our analysis revealed that dynamically changing the geometric relationship of features significantly increased their interpretability for MEG data compared to a fixed orthogonal structure (both fixed orthogonal model and dynamic model peaking at ∼137ms; mean correlation at peak ± s.e.m across 15 subjects: fixed orthogonal model, 0.2532 ± 0.0244; dynamic model, 0.3784 ± 0.0466; Fig. 5a; the responses of VTC peaking at ∼137ms, Supplementary Fig. 8a). This improvement could be attributed to the fitting of noise within MEG signals. To control noise interference, we used the same method to fit the MEG signal with a random matrix. Compared to the improvements provided by the random fitting, adjusting the relationship between the three features significantly improved the interpretation of the MEG data during the period of 110ms to 150ms (one-sided cluster-FEW corrected signed-rank test across 15 subjects for improved correlation of dynamic model > random fitting, p<0.05; Fig. 5a), aligning with the time of visual processing in VTC (Cichy et al., 2014; DiCarlo et al., 2012). Notably, we observed that during baseline, the time before the stimulus presentation, the geometric relationship between features varied within a certain range. However, following stimulus onset, this variation shifted directionally, reaching the peak at ∼135ms, where it exceeded three times the standard deviation observed during baseline (Fig. 5b∼c; Supplementary Fig. 8a). This directed change underscores a correlation between the rotational transformation of features in object recognition and the recognition process itself, indicating an adaptation in feature structure tailored to the need of specific object recognition.

**Figure. 5.**
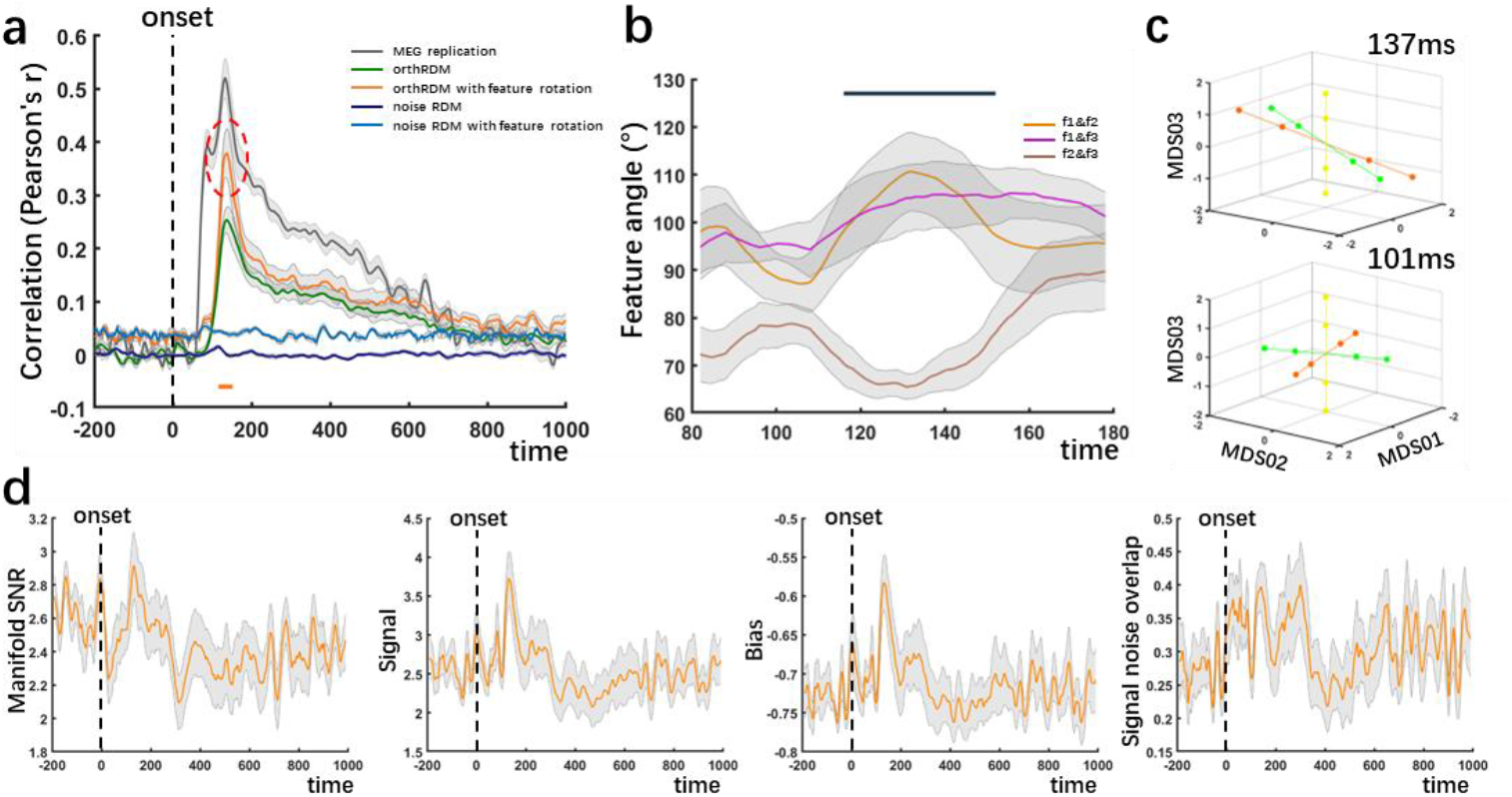
Dynamic of feature structures in object recognition. **a**. The Pearson’s correlation between the MEG RDMs and model RDMs is shown. The green line shows the correlation between the MEG and the fixed orthogonal model of three domain-general features, and the dark blue line shows the correlation between the MEG and the random matrix. The correspondence between the MEG RDMs and the cross-validated fitted feature geometric structure of the two model RDMs are also shown (orange line for the dynamic feature structure and light blue line for the random fitting). The analyses were done separately for each subject, and the results were averaged. Shaded regions indicate s.e.m. across 15 subjects. Significant time points that the dynamic model shows a higher improved correlation than the random fitting model are indicated by thick orange lines at the bottom. (one-sided cluster-FEW corrected signed-rank test across 15 subjects for improved correlation of dynamic model > random fitting, p < 0.05). The gray line indicates the amount of replicable structure in the MEG RDMs across subjects (split-half correlation). Black dots lines indicate the onset of image presentation. **b**. The geometric relationship between three features changes during cognitive processing. The feature angle obtained was reduced noise interference by applying a 20ms moving average window. The thick dark line at the top indicates a significant time point in **a**. Shaded regions indicate s.e.m. across 15 subjects. **c**. The geometric structures of three domain-general features at 100ms and 137ms after stimulus presentation were reconstructed in a low-dimensional space. Each point is a feature node, with colors denoting different domain-general features. **d**. The dynamics of geometric properties of scene manifolds during object recognition. The orange line indicates the group mean, and shaded regions indicate s.e.m. across 15 subjects. Black dots lines indicate the onset of image presentation.

We further investigated how feature dynamic transformation affected the separability of two object manifolds: faces and scenes, given that the majority of stimuli in the dataset fall into these two categories (Supplementary Fig. 8b shows the stimuli of these two categories used in the analysis). After onset, the separability between faces and scene manifolds gradually increased, and reached its peak near 132ms, but this manifold SNR was only slightly higher than the mean during the baseline period without statistical significance (Fig. 5d, manifold SNR; one-sided signed-rank test across 15 subjects for the manifold SNR of baseline < ∼132ms, p = 0.1514, ∼101ms < ∼132ms, p = 0.0473; Supplementary Fig. 8d, manifold SNR). Nonetheless, the detailed analysis of the manifold geometry showed that the signals between manifolds increased significantly following the change in feature structures (Fig. 5d, manifold signal; one-sided signed-rank test across 15 subjects for the manifold signal of baseline < ∼132ms, p<0.01; ∼101ms < ∼132ms, p<0.01, Supplementary Fig. 8d, manifold signal). Moreover, the size of the scene manifold relative to the face manifold significantly increased (Fig. 5d, manifold bias; one-sided signed-rank test across 15 subjects for the manifold bias of baseline < ∼132ms, p<0.01; ∼101ms < ∼132ms, p<0.01, Supplementary Fig. 8d, manifold bias). This change in manifold shape also resulted in a dimension expansion of the scene manifold and a dimension compression of the face manifold (Supplementary Fig. 8c∼d). Overall, the dynamical geometry on manifolds supported the idea that the transformation in feature structure during object recognition was divergent from the mere random fluctuations observed during baseline periods, instead, it embodied a directed rotation, aligning precisely with the demands of specific object recognition tasks. This directional transformation highlights the application of structural knowledge to adapt to various object recognition challenges, providing valuable insights into the underlying computational mechanisms of visual cognition.

## Discussion

Recent studies have suggested that domain-general features in the VTC are critical in understanding how the brain processes and represents object identity (Bao et al., 2020; Vinken et al., 2023). However, how these features collaborate to influence the perception and categorization of stimuli remains unknown. In this paper, we investigate the geometry of domain-general features within the VTC and explore its implications for the separability of object manifolds, which are critical for distinguishing ecologically important objects. Additionally, we propose that the VTC may exhibit dynamic modulation of feature geometry in response to varying object recognition, a computational scheme that could provide new insights into the adaptability of neural processing in visual recognition.

Consistent with our expectation, the fMRI response patterns of nodes on the domain-general features suggested that the features distilled from the ANN were also represented in the VTC, but formed a special geometric structure divergent to the ANN. This geometric difference provided potential biases for VTCs in distinguishing different categories by affecting the geometric properties of object manifolds. By exploring the dynamic nature of these geometric relationships, we found that VTC exhibits dynamic modulation of feature geometry in response to varying object recognition, providing a deeper insight into the adaptability of the visual system and the neural underpinnings of visual recognition.

### Basic domain-general feature structures

Prior work has highlighted the influence of prolonged exposure to specific visual stimuli in the emergence of domain-specific areas in the VTC (Gomez et al., 2019; Srihasam et al., 2014). Conversely, the absence of experience for a particular category would impede the development of corresponding domain-specific areas (Arcaro et al., 2017). The emergence of domain-specific areas shows how the VTC acquires, organizes, and represents domain-general features from an environment, establishing a hierarchical knowledge structure that can be meaningfully applied to any object distinction. Correspondingly, by simulating various visual experiences, ANN models indicate that the statistical structure of an environment influences the formation of structured semantic knowledge in the VTC (Saxe et al., 2019).

To further understand how the VTC feature structure affects inferences of object identity, we conducted experiments with well-controlled visual stimuli to map the three domain-general features distilled through the ANN onto the VTC. Our experiments have revealed that these features are shared between ANN and VTC, but a special geometric relationship between VTC features formed at the individual level, which is divergent from the ANN. Previous studies have reported differences in feature representation between artificial neural networks and the brain (Jiahui et al., 2021; Xu and Vaziri-Pashkam, 2021), such as global preferences (Baker et al., 2020) or lighting (Chang et al., 2021). Our research results indicate that there are also differences in feature organization between the two. We consider that this unique feature structure in VTC reflects the statistical relationships of domain-general features in the visual experience of the participants, potentially benefiting the recognition of some objects, referred to as the basic structure of VTC features. The basic structure of VTC features is not directly tied to particular tasks but rather represents rich semantic knowledge obtained from complex environmental experiences to support identity inference and generalize onto new items and properties. Supporting this idea, we found that modifying the basic structure of features results in changes in the separability of object manifolds. The geometrical properties of manifolds reveal the underlying mechanism of feature structure, affecting separability by constraining the shape of their manifolds and providing a specific predominance for distinguishing objects. Together, these results provide insight into the ways in which long-term visual experiences shape our visual representations and object recognition processes.

A limitation of our conclusion is the indirect relationship between individual visual experience and the basic feature structure of VTC. Due to difficulties in quantifying participants’ visual experiences, we infer our conclusions from the considerable differences between the visual experiences of ANN models and participants and their divergent feature structures. Although ANN is currently the best model for the ventral visual pathway, its model architecture is not completely equal to the ventral visual processing stream, for example, it is difficult to capture the recurrent processing of VVS (Kar and DiCarlo, 2021; Kietzmann et al., 2019). This difference in computational architecture may also be one of the reasons for the divergence of feature structures. In addition, recent studies have shown that the task of training ANNs can impact the structure of features (Dubreuil et al., 2022). Future research is needed to further explore the relationship between the diversity of visual experiences and the basic structure of VTC features. For example, future studies could investigate the VTC feature structure of subjects with special visual experience (Gomez et al., 2019; Ratan Murty et al., 2020; Wang et al., 2020) and elucidate how their experience affects feature structure to provide an advantage in recognizing specific objects.

### Feature rotation and object recognition

The influence of VTC feature structure on object identity inference suggests that changes in feature structure may be involved in the process of object cognition. This idea aligns with the understanding that the visual cortex is not just a static stimulus-response mapping, but is instead influenced by the ongoing task (Hebart et al., 2018; Kay et al., 2023), as supported by our observations while analyzing brain activity during image viewing of participants.

Before the onset of stimuli presenting, we found that the geometric relationship between features oscillated within a certain range. One interpretation of this observation, combined with our previous results, is that the neural activity at this time encodes general information about the relationship between domain-general features in VTC, the basic feature structure. The spontaneous neural activity before the actual viewing of the objects does indeed encode an abstract structure that supports the completion of subsequent tasks (Dragoi and Tonegawa, 2011; Liu et al., 2019). However, due to the absence of rigorous control design in the current dataset, coupled with the pervasive influence of noise inherent in MEG signals, this explanation needs to be treated with caution. Further investigation into the dynamic changes of VTC feature structures in spontaneous neural activity is crucial for gaining insights into how the basic feature structure of VTC plays a role in wildly cognitive tasks.

After the onset, domain-general features rotate in a specific direction, forming a task-specific structure. This effectively enhances the discrimination between related object manifolds to improve separability for downstream read-out. Humans prioritize different aspects of complex stimuli based on their behavioral goals through attention mechanisms (Çukur et al., 2013; Ghose and Maunsell, 2002), which improve the distinguishability of response patterns along task-relevant features while collapsing the task-irrelevant features (Nastase et al., 2017). Attention-based selective enhancement is a potential mechanism that leads to observed feature rotation. Beyond attention, neuron tuning changes are also induced by tasks (Kay and Yeatman, 2017; Popovkina and Pasupathy, 2022). One recent study showed a shift in IT neuron tuning from domain-general to domain-specific features, adapting to the transformation of task goals from face detection to discrimination (Shi et al., 2023). Given that their study used Alexnet to extract both domain-general and -specific features, the rotation of domain-general features might reflect the shifting of features for different task goals. One potential scheme would be feature rotation as a key component of VTC cortical computation, leading to neural tuning features suitable for discrimination tasks within the specific domain. Nonetheless, our findings extend the current computational framework for object recognition, providing insights into how structured knowledge has a dynamic impact on the process of recognizing a single object.

## Conclusion

Our study highlights the significant differences in feature representations between the VTC and ANNs, offering insights into how the brain organizes and utilizes multiple features for object recognition. The adaptable feature structure of the VTC enhances our understanding of the neural basis for organizing knowledge from the environment and applying it flexibly for object identification. These results not only expand our understanding of visual processing but also have implications for the development of brain-inspired neural networks, leading to a deeper exploration of how visual experiences support flexible object recognition.

## Conflict of interest

The authors declare that they have no conflict of interest.

## Acknowledgment

We would like to thank Radoslaw M. Cichy and Aude Oliva for access to the spatiotemporal dynamic data for naturalistic images and Kendrick Kay for access to the Natural Scenes Dataset. This work was supported in part by the National Natural Science Foundation of China under Grant 62020106015 and Grant 61976209.

## Author contributions

Bincheng Wen, Chuncheng Zhang, Changde Du, Le Chang, and Huiguang He conceived and designed the study. Bincheng Wen and Chuncheng Zhang collected human fMRI data. Bincheng Wen analyzed the data and built the computational models. Le Chang and Huiguang He supervised the project. Bincheng Wen wrote the manuscript.

## RESOURCE AVAILABILITY

### Lead Contact

Further information and requests for resources should be directed to and will be fulfilled by the Lead Contact, Huiguang He.

### Materials Availability

This study did not generate new unique reagents.

### Data and Code Availability

The data and code that support the findings of this study are available upon request from the corresponding author by e-mail.

## METHODS

### Participants

We recruited nine healthy participants from the Beijing Normal University (3 males and 6 females, age 21.7 ± 0.6). Five in nine completed all experiments (2 males and 3 females, age 21.4 ± 0.7). All experimental protocols were approved by the Biomedical Research Ethics Committee of the Institute of Automation, Chinese Academy of Sciences and all participants gave their informed consent on the first day of the experiment.

### Visual stimuli and experimental design

#### Domain-general feature mapping

We first randomly selected 200k naturalistic images from ImageNet, spanning 1000 object classes, with 200 images randomly selected from each class (Fig.1a). Then, we extracted the principal components (PCs) as candidate domain-general features by passing all images through Alexnet (Krizhevsky et al., 2012) and conducting principal component analysis (PCA) on the elicited responses of units in the penultimate layer (Fig.1b). Additionally, we refined the selection by focusing solely on the minimum features associated with these three well-known VTC representations: animacy, curvature, and real-world size. When the first three PCs were used to construct orthogonal object feature space (Fig. 1c), the three VTC representations were satisfied at the same time (Supplementary Fig.2), so they were regarded as the candidate domain-general features of VTC.

For each domain-general feature, 200 images with the smallest angle and 200 images with the largest angle were selected from the 200k image set according to the angle between the image coordinates and the feature axis (0°≤angle≤180°), termed “positive” and “negative” representative images of the corresponding feature, respectively. Then, according to the image projection position on the axis, these representative images are further selected and grouped into four feature node subsets, each subset with 50 images, in which two subsets are on the positive end of the feature axis and two subsets on the negative end (Supplementary Fig.1).

In each run of feature mapping fMRI experiments, the positive and negative feature node image blocks of a single domain-general feature were presented alternately. 50 images were randomly selected as control images and presented in all fMRI sessions to serve as a baseline. An example block sequence is shown in Supplementary figure 3a. Each image had a size of 10°×10° visual angles and was displayed for 500ms. Each run consisted of twenty 24s blocks and lasted 480s in the feature mapping experiments. During the fMRI experiments, the participants were instructed to keep their eyes at the central fixation point (0.2° × 0.2°) throughout the experiment and report when the point’s color changed to keep their alertness (which randomly happened 6 times in a block, Supplementary Fig. 3b). For each domain-general feature, 8 runs were collected for each human subject used in the analysis.

#### Meridian mapping

The meridian mapping experiment contained two types of blocks: horizontal and vertical meridians. The stimuli were wedges of a black-and-white checkerboard (flickering at 1 Hz) radiating out from the fixation spot along the vertical and horizontal meridians, occupying 60° and 30° visual angles respectively (Supplementary Fig.3c). Similar stimuli were used previously to determine the vertical and horizontal meridians that define retinotopic visual areas (Lafer-Sousa and Conway, 2013). Two types of blocks were presented in alternation. Each run lasted 480s.

### fMRI data acquisition

The functional and anatomical MRI data were acquired with a Siemens 3T Prisma scanner with a 20-channel head coil. Whole-brain fMRI data were collected using a gradient-echo echo-planar imaging (EPI) sequence (TR = 2,000 ms; TE = 30 ms; flip angle = 90°; slices = 58; imaging matrix = 80 × 80; field of view = 160 mm × 160 mm; 2.5 mm × 2.5 mm in-plane resolution; slice thickness = 2.5 mm). For each session, 10–12 runs were acquired, and each run consisted of 204 functional volumes. A pair of gradient echo images (echo time: 4.92 and 7.38 ms) with the same orientation and resolution as EPI images were acquired to generate a field map for distortion correction of EPI images. High-resolution T1-weighted anatomical images were acquired using a MPRAGE sequence (TR = 2, 530ms; TE = 2.27 ms; flip angle = 7°; acquisition voxel size = 1 mm × 1 mm × 1 mm; 208 sagittal slices). Four or five whole-brain anatomical volumes were acquired and further averaged for better brain segmentation.

### Temporal dynamic data for naturalistic images

To explore the dynamic of VTC feature structure and the constraints of structure on object manifolds at different time points, we used the dataset (Mohsenzadeh et al., 2019) that using MEG and fMRI separately recorded the brain responses during 15 subjects (9 female, 27.87 ± 5.17 years old) passive viewing 156 naturalistic images from the LaMem image set. The majority of stimuli in the dataset fall into two categories: faces and scenes (Supplementary Fig. 8b). Images were presented for 500 ms in randomized order with 1∼1.2s stimulus onset asynchrony (SOA) in MEG experiments. The participants were instructed to press a button and make an eye blink upon detection of a specific eye image.

### Data analysis

#### fMRI preprocessing and general linear modeling

Pre-processing was conducted in MATLAB with SPM12 and custom scripts. The first block of functional images from each run was excluded from the analysis to reach stable magnetization. Data preprocessing included slice time correction, static and dynamic distortion correction, and motion correction of each functional run. The data from all other measurement sessions were co-registered and re-sampled to the same space as the first measurement session data. All analyses were performed in native space; no space normalization was applied. Standard generalized linear modeling (GLM) for the feature nodes of each domain-general feature, the functional localizer, and the meridian mapping was performed on MATLAB and included one regressor per stimulus condition, as well as nuisance regressors for linear drift removal and motion correction per run. The responses of the feature nodes were normalized by the response to the control block to facilitate direct comparison across the domain-general features.

#### Cortical surface reconstruction and segmentation

After the skull was stripped from the average structural image with FSL’s Brain Extraction Tool (University of Oxford), the surface was reconstructed on FreeSurfer. The segmentation was further refined manually. The temporal and occipital lobe region with significant visual responses (p<0.001, paired t-test, control block vs. gray-screen in the feature mapping experiment) was cut out and flattened. The result of meridian mapping was projected onto the surface to divide V1∼V4. The border between V4 and the temporal lobe and anatomical markers including the occipitotemporal sulcus (OTS), posterior transverse collateral sulcus (ptCoS), and parahippocampal gyrus (PHG), was used to help divide VTC. The segmentation results on the flat brain surface were projected onto the native space of each subject as cortical labels of VVS through the mri_label2vol function from FreeSurfer.

#### Data reliability and noise ceiling

To estimate the data reliability as the number of repetitions increases, we randomly split the pooled data into two groups (and repeated this randomization 50 times). The reliability of pooled data for every subject was estimated by first computing the beta estimates per feature node separately for each half of the data and then measuring the Spearman-Brown corrected split-half correlation (repeating this entire procedure 50 times for each binary split) (Supplementary Fig. 4a).

To fairly compare ANN with each brain area, we estimated the noise ceiling of each brain area, which indicates the maximum attainable similarity of the models given the reliability of the recorded data themselves. The lower and upper bounds of the noise ceiling of the fMRI data were calculated following the procedure described by (Nili et al., 2014). Specifically, the upper bound of the noise ceiling for a brain area was established by taking the average of the correlations between each participant’s RDM and the group average RDM including all participants, whereas the lower bound of the noise ceiling for a brain area was established by taking the average of the correlations between each participant’s RDM and the group average RDM excluding that participant (Supplementary Fig. 4b).

#### Participation Ratio (PR) and Explained Variance Dimension

To address how high dimensional trial averaged neural space could be, we computed neural dimensionality, as measured by its participation ratio (Peiran et al., 2017), following:

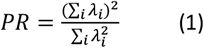

Where *λ*_*i*_ is the *i*^*th*^ eigenvalue of the covariance of the data, measuring the active dimensionality of the eigenspectrum. For VTC dimension estimation, the response of each voxel to each of the four feature nodes at each domain-general feature was normalized by subtracting the mean and dividing by the standard deviation, before the PR computation.

For ANN dimension estimation, the responses of the penultimate layer to the stimuli of all feature node subsets were first extracted. The average response for each feature node subset was regarded as the corresponding activation to the feature nodes. For a fair comparison, we added comparable Poisson noise to the activation of ANN before the PR estimate according to the lower bounds of the VTC responses noise ceiling. Explained variance dimension is defined as the number of principal components required to explain a fixed percentage of the total variance.

#### Representational similarity analysis

##### Representational similarity analysis of fMRI

To compare the geometry of the three domain-general features between VVS and ANN, we compared the RDMs of the feature nodes in each area of VVS and the penultimate layer of the ANN. For V1 ∼ VTC, the multivoxel patterns were first extracted for each of the four feature nodes at each domain-general feature, yielding 12 conditions. Next, the correlation distance (1-Pearson) between neural patterns within these voxels was computed for creating the feature node RDM. For ANN, the feature node RDM was created by Euclidean distances with the average response of the penultimate layer to each feature node subset. The similarity between the feature node RDMs was assessed by taking a Spearman correlation between different feature node RDMs.

##### fMRI and MEG RSA with parameterized model

In order to obtain fine-grained estimates of the geometrical relationship between domain-general features, we also fit a parameterized model to VTCs. We constructed a space of model RDMs by varying three parameters, each controlling the angle between domain-general feature pairs. Let the matrix of 12 feature nodes in the original orthogonal 3-D domain-general feature space be:

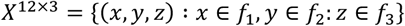

This matrix consists of 3 domain-general features, each aligning with a PC of the penultimate layer of ANN, leading to their mutual orthogonality. The rotation parameter determines the extent to which the representation of the feature is rotated into the frame of reference of the other two features:

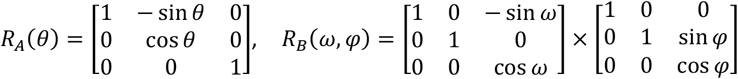

These two rotations were applied to X in a subsequent step, so that the full model was given by:

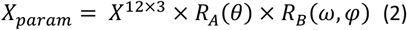

We fit RDMs derived from this model to neural RDMs using a constrained optimization procedure (fmincon in MATLAB) with a least-squares cost function. As the procedure is sensitive to the choice of starting values, we selected the best-fitting parameter values from 180 repetitions with random starting values in fMRI data, and 30 repetitions in MEG data for saving computational resources. The best-fitting RDM was used to visualize the geometrical structure of the domain-general features via projection into three dimensions with classical Multi-Dimensional Scaling (MDS).

##### fMRI RSA for comparison of feature geometry

To compare the geometry of the domain-general features in brains, the feature nodes RDM of the brain were fitted by the RDMs encoded in different models. The first model encoded the orthogonal structure. Since the domain-general features were selected from the PCs of ANN, they formed an orthogonal subspace in the ANN feature space, and the orthogonal structure was encoded by the RDM of the 12 feature nodes in the subspace, we referred to this RDM as orthRDM. This RDM was constructed by computing all pairwise Euclidean distances between the feature nodes in the 3D domain-general feature space. The second model was constructed to encode the mean geometry of domain-general features in VTC. We first averaged the VTC feature angle obtained before to identify the rotation matrix and used it to modify the feature node coordinates in the orthogonal feature space. The corresponding RDM was created by computing all pairwise Euclidean distances between the feature nodes in the new coordinates and referred to as mviaRDM (mean VTC intersection angle RDM). For a given ROI, the upper triangular form of orthRDM and mviaRDM was z-scored and entered into a multiple linear regression model to predict the upper triangular form of the neural RDM:

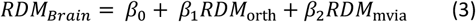

Statistical inference was performed with a group-level t-test of the regression weights. The analyses within ROIs were performed with a leave-one-hemisphere-out cross-validation paradigm. The model RDMs fitted on the rest of the brain hemisphere ROIs were used to predict the RDM of the ROI of the left-out brain hemisphere.

##### Surface-based searchlight RSA

To find brain regions contributing to the formation of VTC-specific structures, we used a model including the RDM encoding the subject’s mean VTC intersection angle to predict brain activity with a surface-based circular searchlight (radius 8 mm) across the whole brain. After projecting the results of each subject to the average brain surface, the statistical inference was performed with a group-level t-test of the regression weights against zero and corrected via a permutation cluster correction based on a cluster forming threshold of p < 0.05.

#### Neural manifold geometric analysis

To explore how the domain-general features structure affects the representation of object information in VTC, we used the geometric properties of the manifolds, including manifold signal, bias, dimensionality, and signal–noise overlap to analyze the manifold SNR (Sorscher et al., 2022). The average error of discriminate pairs of object manifolds (a, b) on test examples of object a is given by *ϵ*_*a*_ = *H*(*SNR*_*a*_), where H(·) is the Gaussian tail function and SNRa is the signal-noise ratio (SNR) for manifold a. The dominant terms of *SNR*_*a*_ are given by:

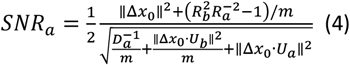

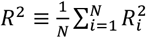 measures the mean squared radius of the manifold along all of the effective dimensions. m represents the sample number of manifold a. The SNR depends on four geometric properties. Signal: 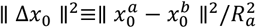 represents the pairwise distance between the manifolds’ centroids, 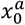 and 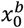. As the pairwise distance between manifolds increases, they become easier to separate, leading to higher SNR. Bias: 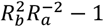 represents the average bias of the linear classifier for a lager manifold. When manifold a is larger than manifold b, the bias term is negative, resulting in a lower SNR for manifold a. Dimension: 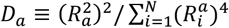 is equal to PR, which quantifies the effective dimension of manifold a. Signal–noise overlap: ∥ Δ*x*_0_ · *U*_*a*_ ∥^2^ and ∥ Δ*x*_0_ · *U*_*b*_ ∥^2^ quantify the overlap between the signal direction Δ*x*_0_ and the manifold axes of variation *U*_*a*_ and *U*_*b*_. The SNR decreases as the overlap between the signal and noise directions increases.

To explore the relationship between VTC feature structures and the separability of object manifolds, we analyzed four ecologically important objects: faces, bodies, words, and places. To minimize the interference caused by variables unrelated to categories, we utilized the stimulus set traditionally used for locating category-selective areas in VTC for analysis (Stigliani et al., 2015). Each category includes two subcategories forming eight objects for manifold analysis: adult faces, baby faces, torso, limbs, numbers, letters, indoor scenes, and architecture. Each object includes 144 images. We first rotated the domain-general features from orthogonal to the previously estimated VTC angle and computed the four geometric properties of the manifold of 8 objects after rotation. The rotation angle of the domain-general features was then used to fit the mean geometric property of 8 object manifolds, respectively. Statistical inference was performed with a group-level t-test of the regression weights with zeros.

To explore how VTC feature structure dynamics affect the separability of object manifolds during object recognition, we examined the manifold SNR of two object categories: faces and scenes, as most stimuli in the MEG dataset belong to these categories. To balance the number of images between categories, we selected 24 images for each category (Supplementary Fig. 3b). We first evaluated the geometrical relationship of domain-general features at each time point, then fine-tuned the features to match the calculated angle, and finally analyzed the four geometric properties and SNR of the two object manifolds.

## Notes

### Competing Interest Statement

The authors have declared no competing interest.

## References

Aparicio, P.L., Issa, E.B., and DiCarlo, J.J. (2016). Neurophysiological Organization of the Middle Face Patch in Macaque Inferior Temporal Cortex. The Journal of neuroscience: the official journal of the Society for Neuroscience 36, 12729–12745.

Arcaro, M.J., Schade, P.F., Vincent, J.L., Ponce, C.R., and Livingstone, M.S. (2017). Seeing faces is necessary for face-domain formation. Nature neuroscience 20, 1404–1412.

Baker, N., Lu, H., Erlikhman, G., and Kellman, P.J. (2020). Local features and global shape information in object classification by deep convolutional neural networks. Vision Research 172, 46–61.

Bao, P., She, L., McGill, M., and Tsao, D.Y. (2020). A map of object space in primate inferotemporal cortex. Nature 583, 103–108.

Bugatus, L., Weiner, K.S., and Grill-Spector, K. (2017). Task alters category representations in prefrontal but not high-level visual cortex. NeuroImage 155, 437–449.

Cadieu, C.F., Hong, H., Yamins, D.L., Pinto, N., Ardila, D., Solomon, E.A., Majaj, N.J., and DiCarlo, J.J. (2014). Deep neural networks rival the representation of primate IT cortex for core visual object recognition. PLoS computational biology 10, e1003963.

Chang, L., Egger, B., Vetter, T., and Tsao, D.Y. (2021). Explaining face representation in the primate brain using different computational models. Current biology: CB 31, 2785-2795.e2784.

Cichy, R.M., Pantazis, D., and Oliva, A. (2014). Resolving human object recognition in space and time. Nature neuroscience 17, 455–462.

Cohen, U., Chung, S., Lee, D.D., and Sompolinsky, H. (2020). Separability and geometry of object manifolds in deep neural networks. Nature communications 11, 746.

Çukur, T., Nishimoto, S., Huth, A.G., and Gallant, J.L. (2013). Attention during natural vision warps semantic representation across the human brain. Nature neuroscience 16, 763–770.

Deng, J., Dong, W., Socher, R., Li, L.J., Kai, L., and Li, F.-F. (2009). ImageNet: A large-scale hierarchical image database. Paper presented at: 2009 IEEE Conference on Computer Vision and Pattern Recognition.

DiCarlo, J.J., and Cox, D.D. (2007). Untangling invariant object recognition. Trends in cognitive sciences 11, 333–341.

DiCarlo, James J., Zoccolan, D., and Rust, Nicole C. (2012). How Does the Brain Solve Visual Object Recognition? Neuron 73, 415–434.

Dragoi, G., and Tonegawa, S. (2011). Preplay of future place cell sequences by hippocampal cellular assemblies. Nature 469, 397–401.

Dubreuil, A., Valente, A., Beiran, M., Mastrogiuseppe, F., and Ostojic, S. (2022). The role of population structure in computations through neural dynamics. Nature neuroscience 25, 783–794.

Epstein, R., and Kanwisher, N. (1998). A cortical representation of the local visual environment. Nature 392, 598–601.

Ghose, G.M., and Maunsell, J.H. (2002). Attentional modulation in visual cortex depends on task timing. Nature 419, 616–620.

Gomez, J., Barnett, M., and Grill-Spector, K. (2019). Extensive childhood experience with Pokémon suggests eccentricity drives organization of visual cortex. Nature human behaviour 3, 611–624.

Grill-Spector, K., and Weiner, K.S. (2014). The functional architecture of the ventral temporal cortex and its role in categorization. Nature reviews Neuroscience 15, 536–548.

Hebart, M.N., Bankson, B.B., Harel, A., Baker, C.I., and Cichy, R.M. (2018). The representational dynamics of task and object processing in humans. eLife 7.

Jiahui, G., Feilong, M., Visconti di Oleggio Castello, M., Nastase, S.A., Haxby, J.V., and Gobbini, M.I. (2021). Not so fast: Limited validity of deep convolutional neural networks as <em>in silico</em> models for human naturalistic face processing. 2021.2011.2017.469009.

Kanwisher, N., McDermott, J., and Chun, M.M. (1997). The fusiform face area: a module in human extrastriate cortex specialized for face perception. The Journal of neuroscience: the official journal of the Society for Neuroscience 17, 4302–4311.

Kar, K., and DiCarlo, J.J. (2021). Fast Recurrent Processing via Ventrolateral Prefrontal Cortex Is Needed by the Primate Ventral Stream for Robust Core Visual Object Recognition. Neuron 109, 164-176.e165.

Kay, K., Bonnen, K., Denison, R.N., Arcaro, M.J., and Barack, D.L. (2023). Tasks and their role in visual neuroscience. Neuron 111, 1697–1713.

Kay, K.N., and Yeatman, J.D. (2017). Bottom-up and top-down computations in word-and face-selective cortex. eLife 6.

Kietzmann, T.C., Spoerer, C.J., Sörensen, L.K.A., Cichy, R.M., Hauk, O., and Kriegeskorte, N. (2019). Recurrence is required to capture the representational dynamics of the human visual system. Proceedings of the National Academy of Sciences of the United States of America 116, 21854–21863.

Konkle, T., and Oliva, A. (2012). A real-world size organization of object responses in occipitotemporal cortex. Neuron 74, 1114–1124.

Kriegeskorte, N., Mur, M., Ruff, D.A., Kiani, R., Bodurka, J., Esteky, H., Tanaka, K., and Bandettini, P.A. (2008). Matching categorical object representations in inferior temporal cortex of man and monkey. Neuron 60, 1126–1141.

Krizhevsky, A., Sutskever, I., and Hinton, G.E. (2012). ImageNet classification with deep convolutional neural networks. In Proceedings of the 25th International Conference on Neural Information Processing Systems - Volume 1 (Lake Tahoe, Nevada: Curran Associates Inc.), pp. 1097–1105.

Lafer-Sousa, R., and Conway, B.R. (2013). Parallel, multi-stage processing of colors, faces and shapes in macaque inferior temporal cortex. Nature neuroscience 16, 1870–1878.

Liu, Y., Dolan, R.J., Kurth-Nelson, Z., and Behrens, T.E.J. (2019). Human Replay Spontaneously Reorganizes Experience. Cell 178, 640-652.e614.

Mohsenzadeh, Y., Mullin, C., Lahner, B., Cichy, R.M., and Oliva, A. (2019). Reliability and Generalizability of Similarity-Based Fusion of MEG and fMRI Data in Human Ventral and Dorsal Visual Streams. Vision (Basel, Switzerland) 3.

Nastase, S.A., Connolly, A.C., Oosterhof, N.N., Halchenko, Y.O., Guntupalli, J.S., Visconti di Oleggio Castello, M., Gors, J., Gobbini, M.I., and Haxby, J.V. (2017). Attention Selectively Reshapes the Geometry of Distributed Semantic Representation. Cerebral cortex (New York, NY: 1991) 27, 4277–4291.

Nili, H., Wingfield, C., Walther, A., Su, L., Marslen-Wilson, W., and Kriegeskorte, N. (2014). A toolbox for representational similarity analysis. PLoS computational biology 10, e1003553.

Pagan, M., Urban, L.S., Wohl, M.P., and Rust, N.C. (2013). Signals in inferotemporal and perirhinal cortex suggest an untangling of visual target information. Nature neuroscience 16, 1132–1139.

Peiran, G., Eric, T., Byron, Y., Gopal, S., Stephen, R., Krishna, S., and Surya, G. (2017). A theory of multineuronal dimensionality, dynamics and measurement. bioRxiv, 214262.

Popovkina, D.V., and Pasupathy, A. (2022). Task Context Modulates Feature-Selective Responses in Area V4. The Journal of neuroscience: the official journal of the Society for Neuroscience 42, 6408–6423.

Ratan Murty, N.A., Teng, S., Beeler, D., Mynick, A., Oliva, A., and Kanwisher, N. (2020). Visual experience is not necessary for the development of face-selectivity in the lateral fusiform gyrus. Proceedings of the National Academy of Sciences of the United States of America 117, 23011–23020.

Rosch, E., Mervis, C.B., Gray, W.D., Johnson, D.M., and Boyes-Braem, P. (1976). Basic objects in natural categories. Cognitive Psychology 8, 382–439.

Saxe, A.M., McClelland, J.L., and Ganguli, S. (2019). A mathematical theory of semantic development in deep neural networks. Proceedings of the National Academy of Sciences of the United States of America 116, 11537–11546.

Sha, L., Haxby, J.V., Abdi, H., Guntupalli, J.S., Oosterhof, N.N., Halchenko, Y.O., and Connolly, A.C. (2015). The animacy continuum in the human ventral vision pathway. Journal of cognitive neuroscience 27, 665–678.

Shi, Y., Bi, D., Hesse, J.K., Lanfranchi, F.F., Chen, S., and Tsao, D.Y. (2023). Rapid, concerted switching of the neural code in inferotemporal cortex. bioRxiv.

Sorscher, B., Ganguli, S., and Sompolinsky, H. (2022). Neural representational geometry underlies few-shot concept learning. Proceedings of the National Academy of Sciences of the United States of America 119, e2200800119.

Srihasam, K., Vincent, J.L., and Livingstone, M.S. (2014). Novel domain formation reveals proto-architecture in inferotemporal cortex. Nature neuroscience 17, 1776–1783.

Stigliani, A., Weiner, K.S., and Grill-Spector, K. (2015). Temporal Processing Capacity in High-Level Visual Cortex Is Domain Specific. The Journal of neuroscience: the official journal of the Society for Neuroscience 35, 12412–12424.

SueYeon, C. (2017). Statistical Mechanics of Neural Processing of Object Manifolds. In eprint arXiv:210600790 (Harvard University, Graduate School of Arts & Sciences.).

Ungerleider, L.G. (1982). Two cortical visual systems. Analysis of visual behavior, 549–586.

Vinken, K., Prince, J.S., Konkle, T., and Livingstone, M.S. (2023). The neural code for “face cells” is not face-specific. Science advances 9, eadg1736.

Wang, X., Men, W., Gao, J., Caramazza, A., and Bi, Y. (2020). Two Forms of Knowledge Representations in the Human Brain. Neuron 107, 383-393.e385.

Wiggett, A.J., Pritchard, I.C., and Downing, P.E. (2009). Animate and inanimate objects in human visual cortex: Evidence for task-independent category effects. Neuropsychologia 47, 3111–3117.

Xu, Y., and Vaziri-Pashkam, M. (2021). Limits to visual representational correspondence between convolutional neural networks and the human brain. Nature communications 12, 2065.

Yao, M., Wen, B., Yang, M., Guo, J., Jiang, H., Feng, C., Cao, Y., He, H., and Chang, L. (2023). High-dimensional topographic organization of visual features in the primate temporal lobe. Nature communications 14, 5931.

Yue, X., Pourladian, I.S., Tootell, R.B., and Ungerleider, L.G. (2014). Curvature-processing network in macaque visual cortex. Proceedings of the National Academy of Sciences of the United States of America 111, E3467–3475.

Yue, X., Robert, S., and Ungerleider, L.G. (2020). Curvature processing in human visual cortical areas. NeuroImage 222, 117295.

